# Sequence and functional characterization of a public HIV-specific antibody clonotype

**DOI:** 10.1101/2021.05.14.444191

**Authors:** Amyn A. Murji, Nagarajan Raju, Juliana S Qin, Haajira Kaldine, Katarzyna Janowska, Emilee Friedman Fechter, Rutendo Mapengo, Cathrine Scheepers, Ian Setliff, Priyamvada Acharya, Lynn Morris, Ivelin S. Georgiev

## Abstract

Public antibody clonotypes shared among multiple individuals have been identified for several pathogens. However, little is known about the limits of what constitutes a public clonotype. Here, we characterize the sequence and functional properties of antibodies from a public HIV-specific clonotype comprising sequences from 3 individuals. Our results showed that antigen specificity for the public antibodies was modulated by the V_H_, but not V_L_, germline gene. Non-native pairing of public heavy and light chains from different donors resulted in antibodies with consistent antigen specificity, suggesting functional complementation of sequences within the public antibody clonotype. The strength of antigen recognition appeared to be dependent on the specific antibody light chain used, but not on other sequence features such as germline or native-antibody sequence identity. Understanding the determinants of antibody clonotype “publicness” can provide insights into the fundamental rules of host-pathogen interactions at the population level, with implications for clonotype-specific vaccine development.

## INTRODUCTION

Antibody discovery from HIV-infected individuals is a hallmark of HIV-1 research, paving the way towards the development of effective therapeutic and vaccine candidates (Bar et al., 2016; Lynch et al., 2015). These discovery efforts have identified broadly neutralizing antibodies (bNAbs) as potential therapeutic candidates and antibodies as templates for engineering antigens to elicit epitope-specific antibody responses to vaccination (Bricault et al., 2019; Jardine et al., 2015; Xu et al., 2018). Large-scale profiling of human antibody repertoires has shown that the antibody response to infection is vast and complex and, therefore, may contain unexplored avenues for vaccine design (Briney et al., 2019; Galson et al., 2015). One currently under-explored area is vaccine design informed by population-level antibody responses (Davis et al., 2019; Kreer et al., 2020). While the majority of a person’s antibody repertoire is unique because of the vast potential diversity generated in part by V (variable), D (diversity), and J (joining) recombination, light chain selection, and somatic hypermutation (SHM) (Briney et al., 2019), individuals can nevertheless possess identical or similar antibodies. Such “public” antibodies have been identified not only for various disease states including HIV-1 infection, SARS-CoV-2 infection, dengue infection, influenza vaccination and others, but also in healthy individuals (Arentz et al., 2012; Ehrhardt et al., 2019; Jackson et al., 2014; Parameswaran et al., 2013; Setliff et al., 2018; Soto et al., 2019; Voss et al., 2021; Yuan et al., 2020). Yet, very little is currently known about the immunological role of public antibodies and their potential as templates for population-level vaccine design.

To gain a better understanding of the properties of public antibodies, we focused on a clonotype that had been previously identified in samples from multiple HIV-infected individuals from the Centre for the AIDS Programme of Research in South Africa (CAPRISA) cohort (Setliff et al., 2018). In particular, this public clonotype included antibodies from three CAPRISA donors, with publicness defined by the same V_H_-gene, J_H_ gene, and junction length, and CDRH3 amino acid sequences of high identity among donors (Setliff et al., 2018). In that study, two antibody clonotype members with natively paired heavy and light chains from the public clonotype were produced experimentally and confirmed to be HIV-specific. Subsequent analysis of the antibody sequencing data revealed the existence of additional antibody sequences with high CDRH3 identity to the antibodies from the public clonotype but different V_H_ and/or V_L_ genes. Therefore, here we sought to build on our previous work (Setliff et al., 2018) by investigating the genetic and phenotypic characteristics that define the members of this public antibody clonotype. The resulting analysis offers insight into the public antibody response in the context of chronic HIV-1 infection and explores the boundaries of antibody “publicness.”

## Results

### Identification of antibodies with high CDRH3 sequence identity from multiple HIV-infected donors

The antibody sequences from the three CAPRISA donors were combined and complete linkage clustering was performed to assign clonotype membership for each sequence (Gupta et al., 2015). In contrast to our previously published work (Setliff et al., 2018), we expanded our parameters such that sequences were clustered using the following criteria: CDRH3 amino acid sequence identity of at least 70% with the same CDRH3 and junction length and no consideration for V_H_- and J_H_-gene usage. This allowed for a more inclusive definition of potential public antibodies that would enable systematic exploration of the boundaries of antibody publicness. Among the 24,218 clonotypes encompassing sequences from one or more of the three donors, clonotype #13905 contained the previously reported public heavy chain sequences (Setliff et al., 2018) and was selected for further analysis. Clonotype #13905 included 171 nucleotide sequences from all three donors (donors CAP314, CAP248 and CAP351), spanning four different V_H_-gene assignments (*IGHV1-69, IGHV5-51, IGHV1-18* and *IGHV3-23*). Since paired heavy-light chain sequences were available for the datasets for donors CAP314 and CAP248, the corresponding light chain sequences were also retrieved. Two different light chain genes were identified in sequences from clonotype #13905: IGKV1-27 in both donors CAP314 (20% of light chain sequences from that donor) and CAP248 (100%), and IGKV3-20 in donor CAP314 (80% of light chain sequences from that donor).

### CDR3 sequences from different donors are not less similar than they are within donors

To determine whether the CDR3 sequences from the same donor exhibited greater levels of similarity compared to sequences from other donors, we constructed Hamming distance matrices among all the unique CDRH3 and CDRL3 sequences, respectively, from clonotype #13905. The 171 sequences in the public clonotype comprised 25 unique CDRH3s, including 16 from donor CAP351, five from donor CAP314, three from donor CAP248 and one shared between donors CAP314 and CAP351. It also included six unique CDRL3s, with four from donor CAP314 and two from donor CAP248 (Figure 1A). The CDRH3 sequence distance values ranged from zero to four, with lower values corresponding to greater identity between two CDRH3 sequences (Figure 1A). High CDR3 similarity was observed both within donors (mean: 2.57, SEM: 0.08) as well as among donors (mean: 2.46, SEM: 0.06), with some CDR3s exhibiting greater similarity among, as opposed to within, donors (Figure 1B).

**Figure 1.**
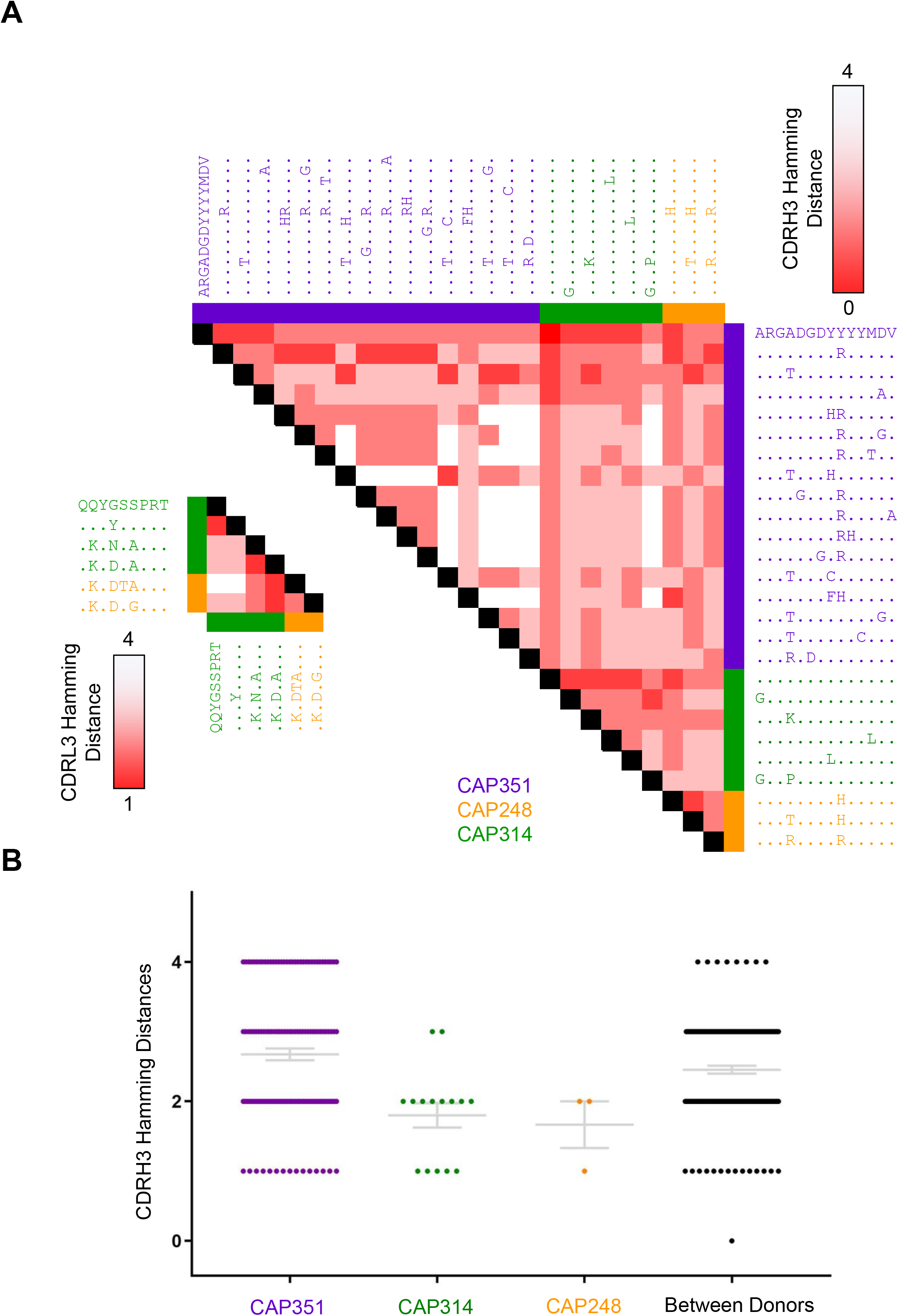
Association of CDR3 Regions Within and Among Members of the Public Clonotype. a. Heatmap of Hamming distance between pairs of unique CDR3s of the sequences in clonotype #13905. The Hamming distance values ranged from 0 (red) to 4 (white) for the CDRH3 and from 1 (red) to 4 (white). For heavy and light chains, the Hamming distance between 25 unique CDRH3 and 6 unique CDRL3 are shown, respectively. Each CDR3 is colored based on the respective donor. An arbitrary sequence was used as a template to visualize amino acid identity (dots) or differences (letters) for the different sequences in the alignment. b. Grouping of Hamming distance values between pairs of unique CDRH3 of the sequences in clonotype #13905 within each donor and among donors with mean with SEM for each group shown in grey.

### Unique CDR3 sequences can be associated with multiple diverse V genes

To investigate the patterns of pairing between CDR3 sequences and germline V genes, we constructed a Hamming distance-based network graph for the antibody sequences from clonotype #13905 (Figure 2). The majority (97%) of heavy chain sequences utilized *IGHV1-69*, but antibodies using three other heavy chain germline genes were also observed: *IGHV1-18* (1.2%), *IGHV3-23* (1.2%), and *IGHV5-51* (0.6%). Of note, the antibodies from donors CAP314 and CAP248 only utilized *IGHV1-69*, whereas donor CAP351 utilized all four of the V_H_-genes. We observed two unique CDRH3 sequences from donor CAP351 that each utilized two different V_H_-genes: ARGADGDYYYYMAV (*IGHV1-69* and *IGHV5-51*) and ARGADGDYRYYMDV (*IGHV1-69* and *IGHV1-18*). These results suggest that the same CDRH3 sequence can be associated with multiple diverse V_H_-genes. With the exception of a single node in the heavy chain portion of the graph, all other antibody sequences from clonotype #13905 were within a Hamming distance of one from at least one other sequence in that clonotype, revealing a tight network of sequence similarity among members of this clonotype, even among different donors (Figure 2). For the light chains, *IGKV1-27* was identified for sequences from both donors CAP248 and CAP314, while donor CAP314 also had antibodies that utilized the *IGKV3-20* germline gene. Overall, a diversity of germline genes was observed for both the heavy and light chains in clonotype #13905, without any clear patterns of association with specific CDR3 sequences.

**Figure 2.**
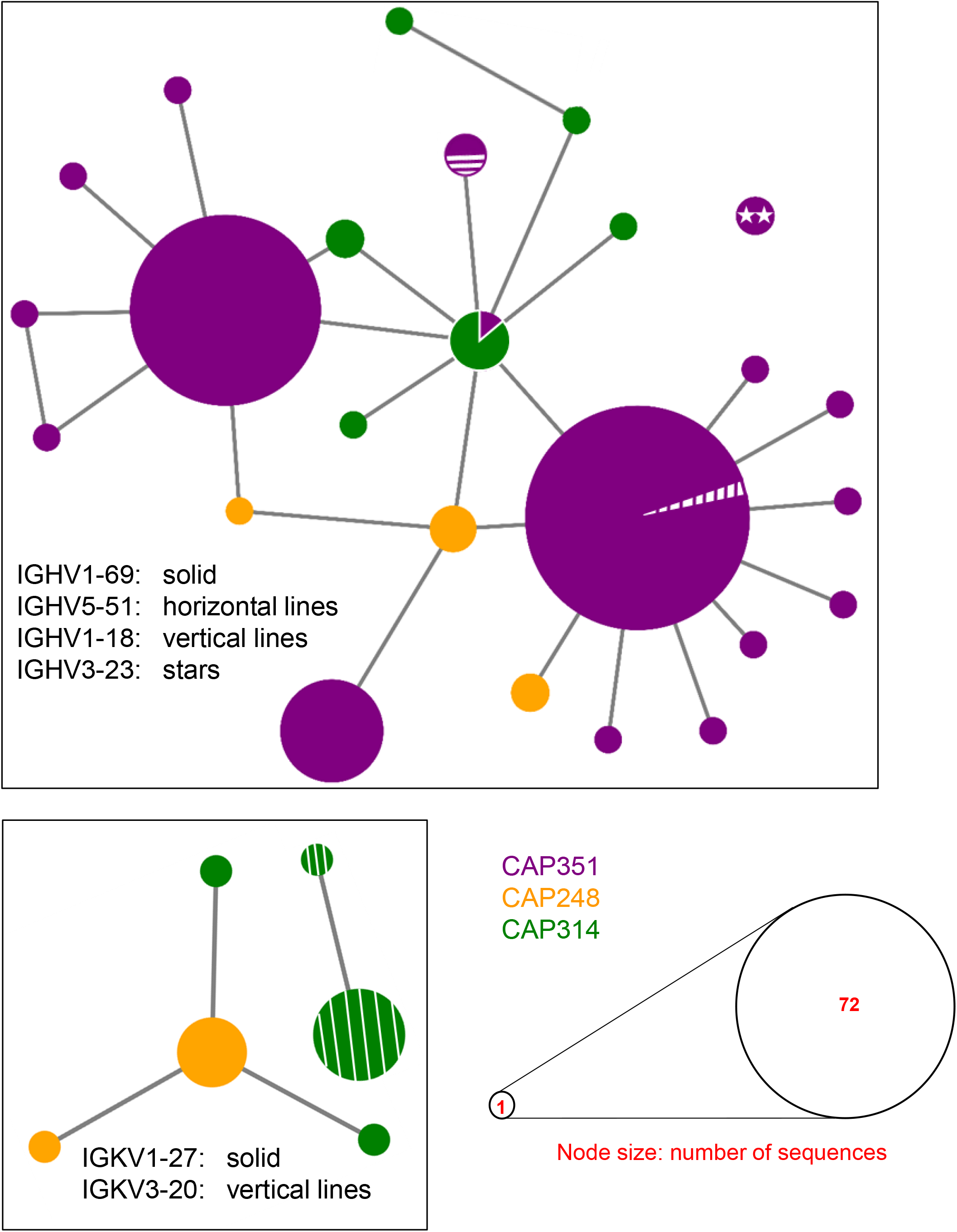
Network Graph Representation of Unique CDR3s in Clonotype #13905. Each node represents a unique CDR3 and the node diameter is proportional to the corresponding number of sequences, which varied from 1 to 72 for heavy chain and 1 to 7 for light chain. Nodes are connected by an edge if their Hamming distance is 1. Node colors correspond to the respective donor for the given CDR3 sequence, with multiple colors in a single node representing a CDR3 that is shared by multiple donors. The node fill pattern corresponds to the V-genes used for the given set of sequences, with multiple patterns in a single node representing different V-genes associated with the same CDR3 sequence.

### Defining the limits of what constitutes a public antibody clonotype

The observation that multiple diverse V_H_ and V_L_ germline genes can be associated with highly similar CDR3 sequences led us to address the limits of what constitutes a public antibody clonotype. To assess this question, we selected a diverse set of antibody heavy and light chain sequences for experimental validation, making sure sequences from all three donors and all observed V_H_ and V_L_ genes were selected. The set included nine heavy chain sequences (six from CAP351, two from CAP314 and one from CAP248) and three light chain sequences (two from CAP314 and one from CAP248). Of these, five, two, one and one sequences were from *IGHV1-69, IGHV3-23, IGHV1-18*, and *IGHV5-51*, respectively; and two and one were from *IGKV1-27* and *IGKV3-20*, respectively. Among the nine heavy chain sequences, three pairs utilized identical CDRH3s (ARGADGDYRYYMDV, ARGADGDYYYYMDV, ARGRDDDYYYYMDV), and interestingly, sequences in one of these pairs (ARGADGDYRYYMDV) utilized different V_H_-genes (*IGHV1-18* and *IGHV1-69*). Thus, the selected set was representative of the diversity of sequences found in clonotype #13905.

All natively paired heavy-light chain sequences, as well as all other non-native heavy-light chain pairs from all three donors (a total of 27 unique antibody heavy-light chain pairs) were successfully expressed as recombinant IgG proteins. Thus, we sought to determine whether and to what extent these antibodies could recognize HIV-1 Env-derived antigens. Two of these antibodies, CAP248_30 and CAP314_30 (native heavy-light chain pairs from donors CAP248 and CAP314), had been previously validated (Setliff et al., 2018) and were also confirmed to be HIV-specific in our experiments (Figure 3, S1). We tested all 27 heavy-light chain pairs against two different HIV Env-derived antigens, clade CRF01_AE 93TH975 gp120 monomer (Figure 3A, S1A) and clade A BG505.SOSIP.664 prefusion-stabilized gp140 trimer (Sanders et al., 2013) (Figure 3A, S1B). Only antibodies with heavy chains utilizing *IGHV1-69* were able to bind the gp120 monomer (Figure 3A, S1A), suggesting the importance of V_H_-gene usage for antigen recognition. Interestingly, the CAP351_04 (*IGHV1-69*) and CAP351_01 (*IGHV1-18*) heavy chains had the same CDRH3 sequence, but different V_H_-genes; however, only CAP351_04 (*IGHV1-69*) bound 93TH975 gp120. In contrast, all IGHV1-69 heavy chains paired with light chains utilizing any of the three V_L_ genes were able to bind gp120 monomer, albeit to different extents (Figure 3A, S1A). Together, these results suggest potential V_H_, but not V_L_, germline gene-mediated antigen specificity for this public clonotype.

**Figure 3.**
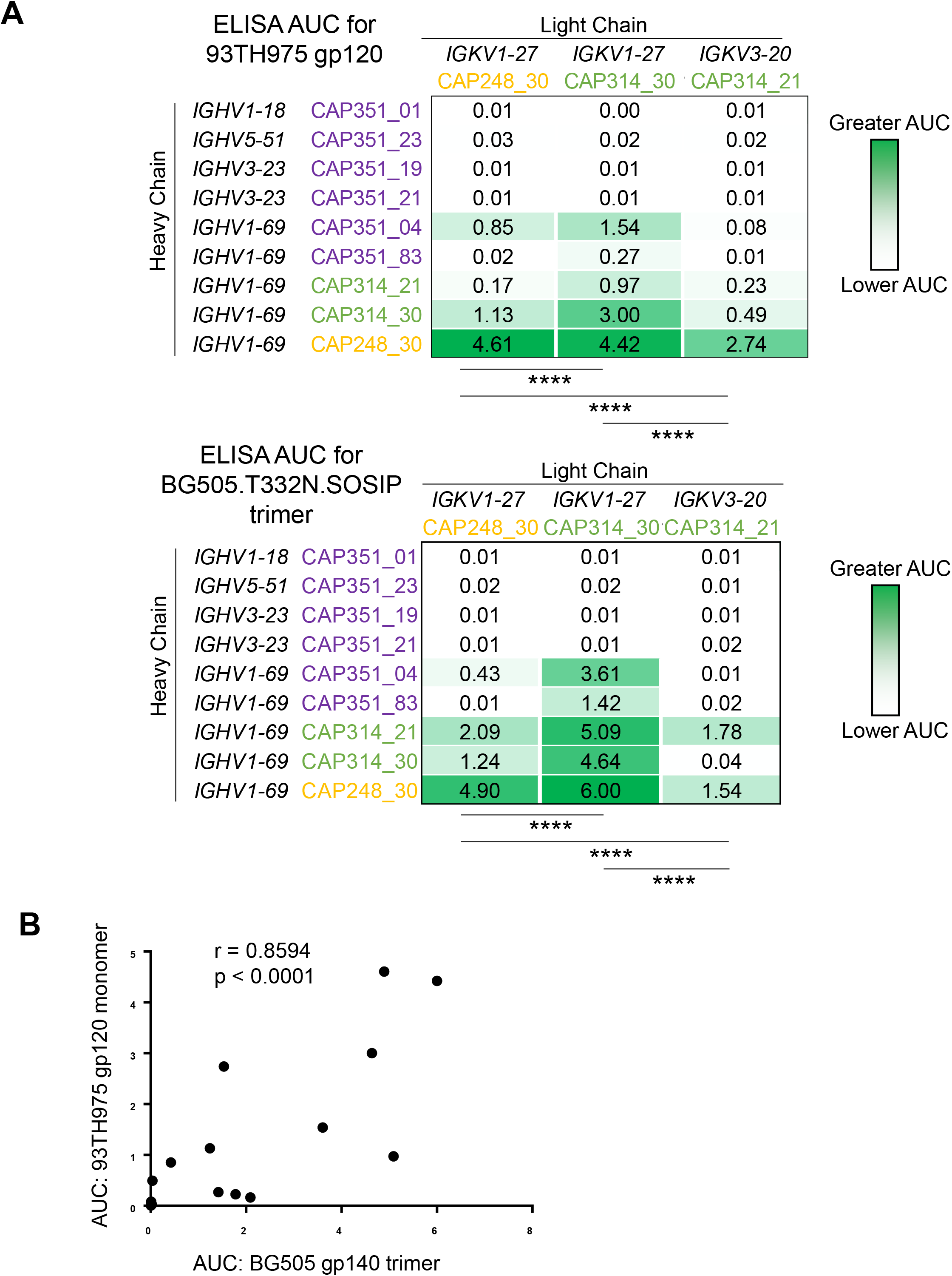
Public Antibody Recognition of HIV-1 Protein. a. Binding data for each antibody (both native and non-native heavy-light chain pairs) against 93TH975 gp120 or BG505.T332N.SOSIP.664 are displayed as a heatmap of AUC analysis calculated from the ELISA curves in Figure S1a. Statistical significance was determined via 2-way ANOVA p-value (p < 0.0001) corrected for multiple comparisons using Tukey’s multiple comparisons test for both monomer and trimer. b. Spearman correlation between antibody binding to HIV-1 monomer (x-axis) and trimer (y-axis).

The strength of binding to the monomer appeared to be associated with the choice of light chain pairing with a given *IGHV1-69* heavy chain (p<0.0001, 2-way ANOVA with p-value corrected for multiple comparisons using Tukey’s multiple comparisons test), suggesting that while there appears to be greater promiscuity for the choice of light chain compared to heavy chain, strength of antigen reactivity can be modulated by optimizing heavy chain-light chain pairings. We next determined if the antibodies from this public clonotype recognized gp140 trimer, which is designed to mimic a neutralization-sensitive prefusion conformation of Env (Sanders et al., 2013) (Figure 3A, S1B). For most antibodies, there was markedly increased binding to BG505.T332N.SOSIP.664 single-chain trimer (Georgiev et al., 2015), though the antibody-antigen binding patterns between monomer and trimer showed a significant correlation (p < 0.0001, Spearman correlation) (Figure 3B). Binding to different forms of the BG505 trimer for the native CAP248_30 and the non-native CAP248_30_H_/CAP314_30_L_ antibody pairs was further validated by surface plasmon resonance (Figure S2).

Overall, the finding that only antibodies using *IGHV1-69* heavy chains were capable of recognizing HIV-1 antigens, despite also testing sequences with highly similar or even identical CDR3 sequences but different V_H_ gene usage, indicated that the public antibody clonotype may be restricted to only sequences with *IGHV1-69* but with either *IGKV1-27* or *IGKV3-20*. This conclusion was further reinforced by the fact that non-native pairs of heavy and light chains from either the same or different donors could successfully recognize HIV-1 Env, indicating functional complementation of antibody sequences from the public clonotype.

### Public antibodies exhibit consistent epitope specificity and virus neutralization

We previously determined that binding to HIV-1 Env by CAP248_30 and CAP314_30 was affected by the receptor binding site (CD4bs) epitope knockout, D368R (Setliff et al., 2018). To confirm whether other members of the public clonotype also mapped to CD4bs, we generated epitope knockouts in the context of the BG505.T332N.SOSIP.664 trimer and confirmed that all tested heavy-light chain pairs were indeed affected by CD4bs knockout mutations (D279K and D368R) (Figure S3). These results suggest that non-native pairings of heavy and light chains from the same or different donors did not affect epitope targeting, further corroborating the conclusion that these antibody sequences which were shared among all three donors, are indeed public. Further, in pseudovirus neutralization assays, both native and non-native heavy-light chain pairs exhibited generally consistent ability to neutralize tier 1 HIV-1 strains (Figure S4), in agreement with our previously published work (Setliff et al., 2018) for the native CAP248_30 and CAP314_30 antibodies.

### Public antibody sequences exhibit high similarity both within and among donors

To visualize the overall sequence similarity among antibodies from different donors, we generated a phylogenetic tree with the sequences from the public antibody clonotype (Figure 4A). Varied levels of somatic hypermutation were observed both in the heavy chains (2.64%-11.16%) and the corresponding light chains (1.4%-4.9% for *IGKV1-27* and 1.04%-5.58% for *IGKV3-20*). Further, multiple clades were observed particularly in the heavy chain tree, suggesting a diversity of antibody evolution within this clonotype (Figure 4A). Of note, however, sequences from different donors were interspersed in the trees. This suggests that in some cases, greater similarity was observed among sequences from different donors, as opposed to sequences from the same donor. This observation of among versus within donor similarities was further supported by the Hamming distances for these sequences (Figure 4B), with distances ranging between 12-22 among donors and 1-19, 2-17, and 6-24 within donors CAP351, CAP248, and CAP314, respectively. Together, these results suggest a diversity of pathways in which antibodies from this clonotype evolved within each donor, although inter-donor similarities were also observed.

**Figure 4.**
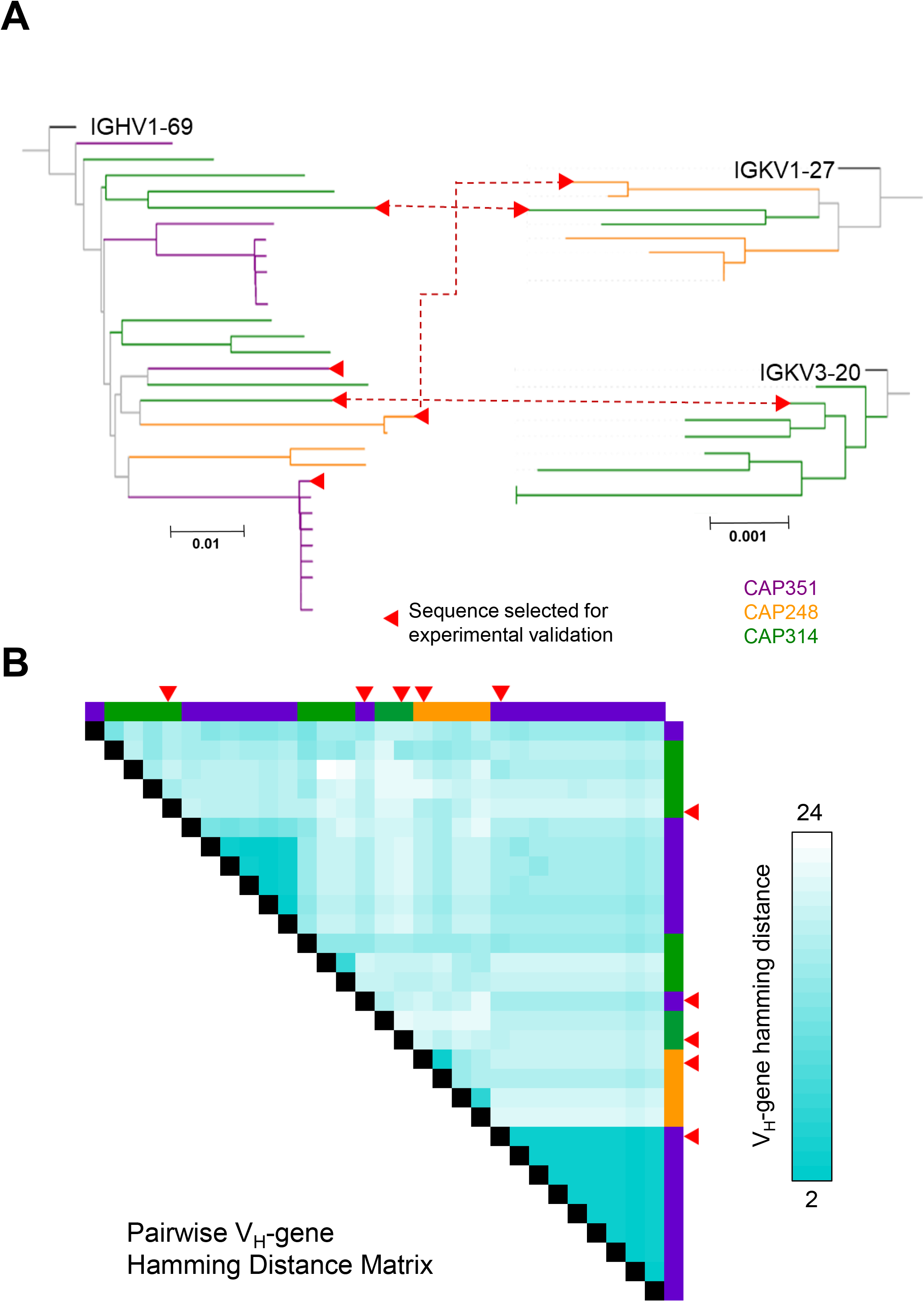
Association of V-Genes Within and Among Members of the Public Clonotype. a. Phylogenetic tree representation of heavy and light chain antibody sequences (colored by donor) in the public clonotype, along with the respective germline genes (black). Dotted lines between heavy and light trees represent natively paired heavy and light chains that were selected for experimental validation. Triangle symbols highlight additional sequences selected for experimental validation. b. Heatmap of Hamming distance values between the V_H_ regions of pairs of sequences (row and columns, colored by donor) in the public clonotype. The Hamming distance values ranged from 2 (cyan) to 24 (white). Triangle symbols highlight sequences selected for experimental validation.

### Public antibodies include distinctive somatic hypermutation changes

To better understand the types of somatic hypermutation changes that are characteristic of the public antibodies, we analyzed the per-residue frequency of mutations from germline (Figure S5A). While in each donor, a large number of residue positions retained their germline identity in the majority of sequences, a number of residue positions had high frequency of mutations compared to germline (Figure S5A). Of note, several of these residue positions overlapped in multiple donors, with the levels of somatic hypermutation and residue entropy at each position having a significant correlation between all three donors (Spearman correlation test with p-value correction for multiple comparisons using Benjamini-Hochberg method) (Benjamini and Hochberg, 1995). This finding suggests the existence of somatic hypermutation hotspots that are common to all three donors (Figures S5A, B).

To interrogate whether the somatic hypermutation hotspots in the *IGHV1-69* antibodies are characteristic of the public antibody clonotype, we compared them to a set of unrelated representative antibody sequences that also utilized the *IGHV1-69* germline gene, retrieved from cAb-Rep (Curated Antibody Repertories) (Guo et al., 2019). Although for the majority of residue positions, no major differences among the public antibody sequences and the reference dataset were observed, there were also several residue positions with >4-fold higher frequencies of non-germline amino acid identities in the public antibody sequences, suggesting the existence of unique *IGHV1-69* mutations that are specific to the public antibody clonotype (Figures 5A, B and S6).

**Figure 5.**
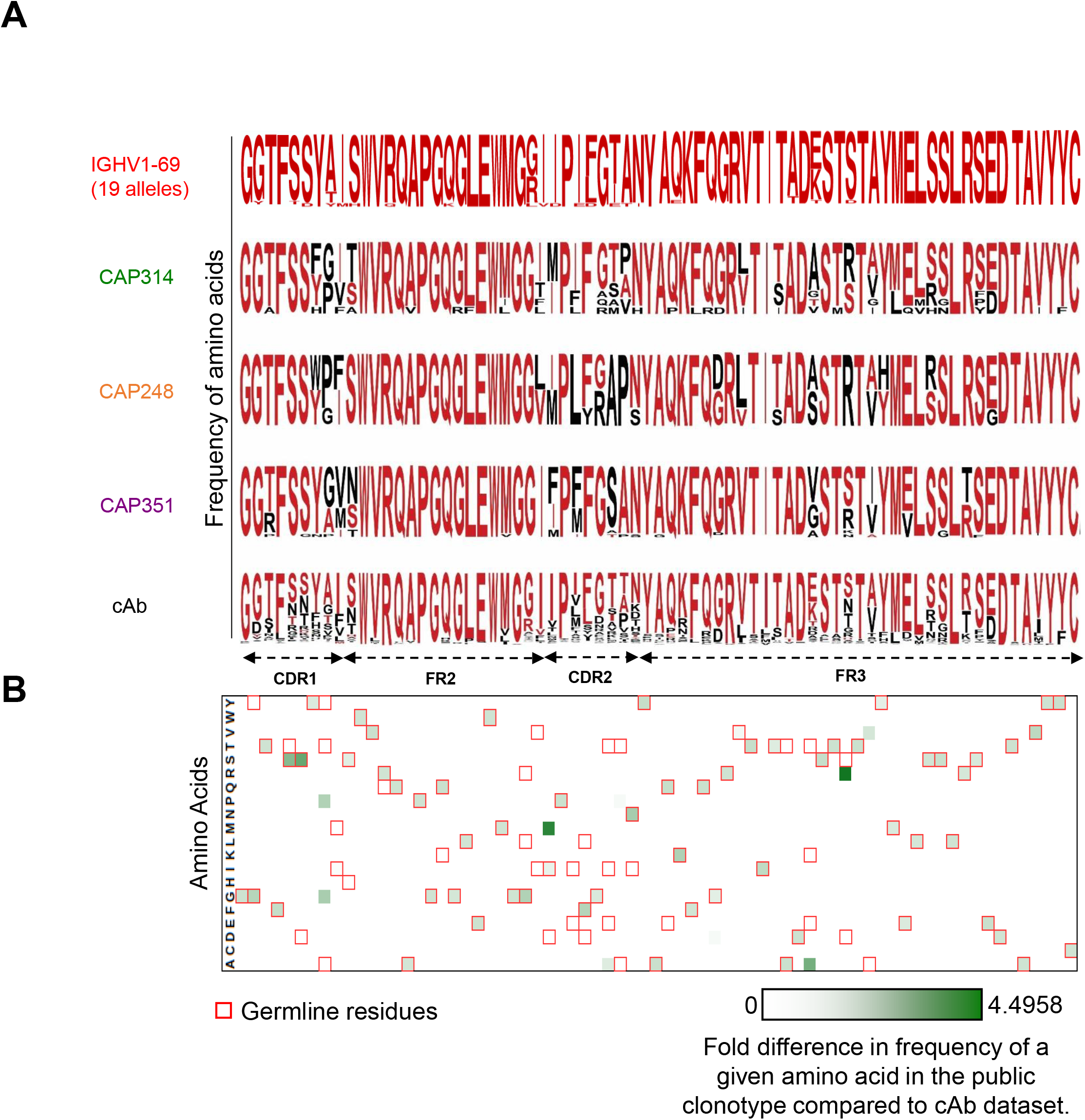
Comparison of Public Antibody Sequences to cAb Dataset. a. Conserved sequence features in the public clonotype. Sequence logo plot representation of *IGHV1-69* germline sequence (generated using sequences of 19 different alleles), heavy chain sequences in the public clonotype and a reference (cAb) dataset of *IGHV1-69*-based antibody sequences. Since the NGS data for donor CAP351 only included sequences between CDR1 and FR3 (Setliff et al., 2018), only that region is shown in the alignment. For each of the three donors and the cAb sequences, residues that are identical to the germline sequences are shown in red, while all other amino acids for a given position in the sequence alignment are shown in black. Height of letters corresponds to the frequency with which the respective amino acid is observed for the given position in the sequence alignment. b. For each residue position in the sequence alignment, the minimum fold difference between the amino acid frequency for a given residue position in the three donors in the public clonotype and cAb dataset is shown as a heatmap, with a color scale of white (0) to green (4.4958). The *IGHV1-69* germline residues (based on sequences of 19 alleles) in each position are highlighted in red boxes.

### Heavy chain identity to either germline or native antibody sequence does not modulate Env trimer recognition

We next examined additional factors that could affect antigen recognition by the different antibody variants from the public clonotype (Figure 3). We first explored the potential role of heavy chain identity to germline sequence. Notably, antibodies at both ends of the V_H_ germline identity scale showed lower levels of HIV-1 protein binding, whereas some of the strongest binding was observed for antibodies with intermediate V_H_ germline identity (Figure 6A). We then explored whether antigen binding could be dependent on the sequence identity of the heavy chain in a given antibody pairing to the heavy chain present in the native pair for the given light chain (Figure 6B). For CAP248_30_L_, the native heavy-light chain pairing was optimal in terms of binding to both gp120 monomer and to the stabilized trimer; however, high identity to native for the non-native heavy chains was not necessarily associated with improved antigen recognition (Figure 6B). Further, the other two light chains, CAP314_30_L_ and CAP314_21_L_, exhibited better antigen recognition when paired with non-native, compared to their native, heavy chains (Figure 6B). Together, these data indicate that heavy chain identity to germline or native sequence may not be strong determinants of antigen recognition by this public antibody clonotype.

**Figure 6.**
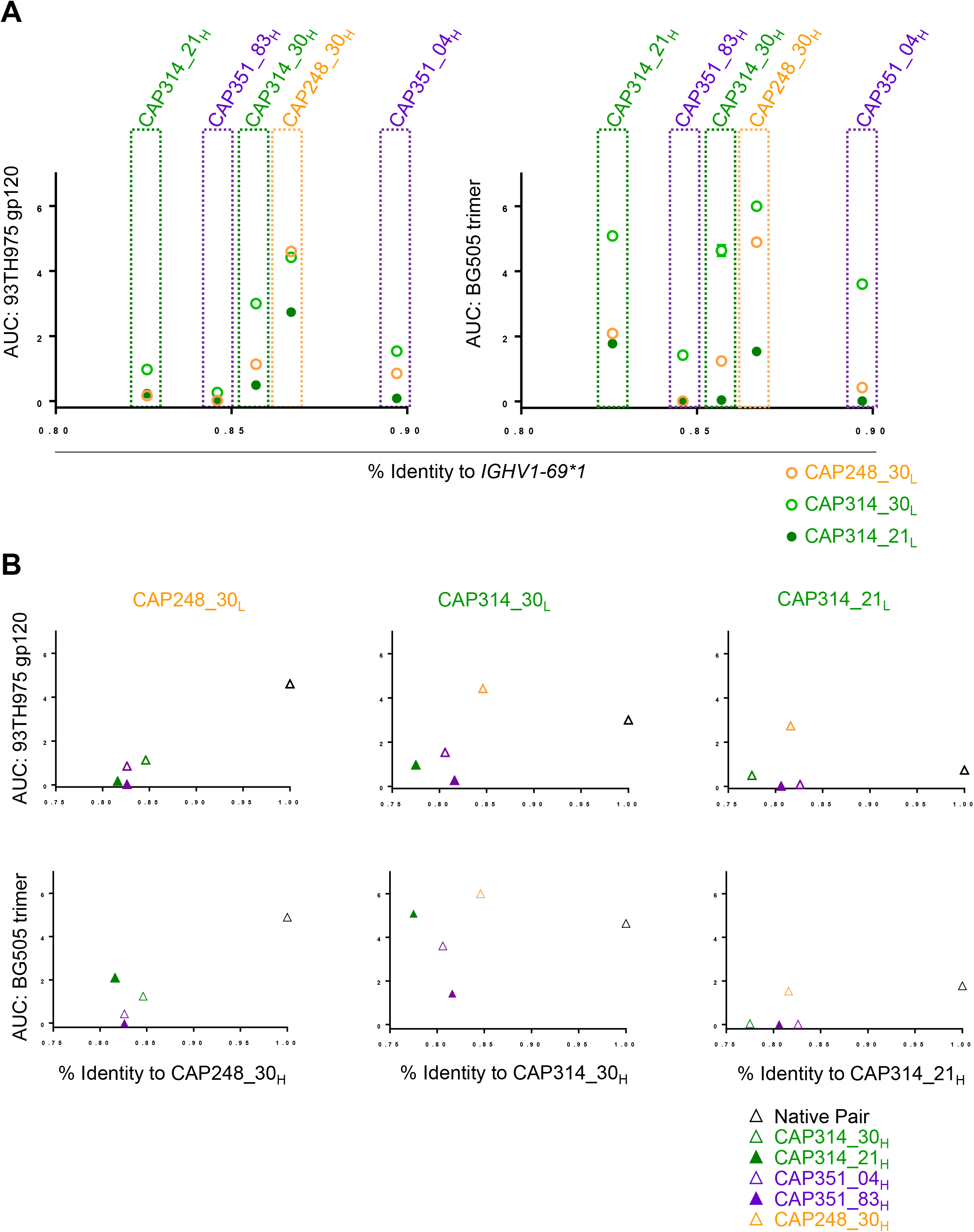
Association Between Antigen Binding and (a) Germline or (b) Native Antibody Sequence Identity. a. Antibody ELISA AUCs (y-axis; left – monomer; right – trimer)) were plotted against percent identity to the germline gene *IGHV1-69*01* (x-axis). Dots correspond to the light chain used in each antibody and dotted boxes group antibodies by the respective heavy chain used. Colors show the source donor; for donor CAP314, the two different light chains are shown as open and closed circles, respectively. b. For each light chain, antibody ELISA AUCs (y-axis; top – monomer; bottom – trimer) were plotted (triangles) against the sequence identity of the corresponding heavy chain to the natively paired heavy chain (x-axis). Each plot corresponds to the titled light chain paired with each heavy chain, denoted by triangles. Colors correspond to source donor, with multiple heavy chains per donor represented with open and closed triangles, respectively.

## DISCUSSION

Public antibody sequences have become an important research direction for a variety of diseases, including dengue, influenza, HIV-1, and SARS-CoV-2 (Parameswaran et al., 2013; Jackson et al., 2014; Setliff et al., 2018; Yuan et al., 2020). Yet, our understanding of what constitutes antibody publicness has been limited to date. At one extreme, stringent definitions that restrict public clonotypes to only identical antibody sequences have been employed (Soto et al., 2019). Such a stringent approach aims to guarantee complete confidence in the identification of truly public antibodies. However, this approach fails to account for the diversification potential of antibodies undergoing evolution in response to antigen exposure. For example, antibodies from an individual clone from a single donor have been identified with up to 50% or more divergence in CDRH3 sequence (Wu et al., 2015). It is therefore important to understand what levels of sequence-based similarity can be reasonably used for defining public clonotypes among antibodies from multiple individuals.

To that end, here we characterized a previously identified public clonotype found in a cohort of HIV-infected donors by expanding selection criteria and performing phenotypic and genotypic analyses to more carefully define the limits of antibody publicness for this clonotype. To achieve this, we explored the choice of V_H_ genes, the choice of V_L_ genes, and a range of CDR3 identities for sequences in the putative clonotype. We also explored the potential of heavy and light chains from different antibodies and different donors to produce functional antibodies with unaltered antigen specificity to show functional complementation between non-native heavy-light chain pairs and further support the phenotypic similarities among different antibodies in this public clonotype. Of course, as expected, the specific choice of heavy and light chain pair appeared to be important for strength of antigen recognition. Whether our findings for the specific clonotype studied here will be generalizable for other public antibodies and in contexts beyond HIV-1 infection will be of significant interest in the antibody field. Additional investigation of public clonotypes may shed light on specific, population-level responses to infection and vaccination. A comprehensive assessment of the publicness of antibody repertoires has the potential to make significant contributions to vaccine development for difficult targets such as HIV-1, influenza, and other diseases.

## Acknowledgements

We thank Tandile Hermanus and Carol Crowther for assistance in characterizing the antibodies described in this manuscript; and members of the Georgiev lab for comments on the manuscript. This work was supported in part by institutional funding from Vanderbilt University Medical Center and: (I.S.G.) NIH R01 AI131722; (A.A.M.) NIAID T32AI112541; (L.M.) the Medical Research Council of South Africa, and NIAID U19 AI51794 (CAPRISA) and 1U01AI136677; (P.A.) R01 AI145687; (ATRECA) Global Health Vaccine Accelerator Platforms ID-48 funded by the Bill and Melinda Gates Foundation. The funders had no role in study design, data collection and analysis, decision to publish, or preparation of the manuscript.

## Materials and Methods

### Reagents

The following reagents were obtained from the AIDS Research and Reference Reagent Program, Division of AIDS (DAIDS), National Institute of Allergy and Infectious Diseases (NIAID), National Institutes of Health (NIH): HIV-193TH975 gp120 from Steve Showalter, Maria Garcia-Moll, and the DAIDS, NIAID; Anti-HIV-1 gp120 Monoclonal (3BNC117) from Dr. Michel C. Nussenzweig (Shingai et al., 2013); Anti-HIV-1 gp120 Monoclonal (VRC01), from Dr. John Mascola (cat# 12033) (Wu et al., 2010).

### Sequencing data analysis and clonal selection

We used preprocessed B-cell repertoire data of donors CAP351, CAP314 and CAP248 from our previously published sequencing data which is available for public access under BioProject PRJNA415492 (Setliff et al., 2018). For donor CAP351, heavy chain variable gene sequences were available from three time points (pre-infection, six months post-infection and three years post-infection) whereas for donors CAP314 and CAP248, paired heavy-light chain sequences were available for a single time point. Preprocessed sequences were annotated using IgBLAST (Ye et al., 2013) to assign gene information compared to the germline repertoire obtained from IMGT (Lefranc et al., 2015). To find the public clonotype, irrespective of V-gene and J-gene identity, we combined data from all three donors and performed clonal clustering using Change-O (Gupta et al., 2015) with the following criteria: complete linkage, same CDRH3 length and 70% CDRH3 amino acid sequence identity.

### Public clonotype analysis

For each of the heavy chain and light chain sequence sets, sequences were grouped based on the respective V-genes. Hamming distances were computed between all the pairs of unique CDR3s. Further, sequences were grouped based on CDR3, and a network graph was generated using each unique CDR3 as a node; an edge between two nodes was shown only for Hamming distance value of one. A multiple sequence alignment with phylogenetic analysis was performed using Clustal Omega (Madeira et al., 2019) by also including the respective germline V-gene sequences. Phylogenetic trees were annotated and visualized using iTol (Letunic and Bork, 2019); for better visualization, sequences were clustered based on 98% sequence identity and one (original and/or consensus) or more sequences (if they came from different donors or V-genes).

Residue-wise analysis was performed by computing somatic hypermutation (SHM) and entropy values. SHM was computed using in-house scripts that compared each sequence in the public clonotype to the respective germline sequence. This revealed the frequency of mutation at each residue position, which can vary from 0 (non-mutated) to 1 (always mutated). Entropy values were calculated by using a log-based formula via Bioedit (Hall et al,1999). The entropy of a residue position represents the frequency of different amino acids at that position. Entropy values at each residue position varied from 0 (only one type of amino acid at that position) to 4.322 (different frequencies of multiple amino acids at that position).

Sequences in the public clonotype were also compared to sequences in the cAb-Rep database (Guo et al.,2019). cAb-Rep is a curated database containing, at the time of analysis, antibody sequences from 306 B cell repertoires from 121 donors. Among the 267.9 million heavy chain sequences from cAb-Rep, sequences with gene *IGHV1-69* were extracted, and sequences with less than 98% identity and SHM >10% were included in the “cAb dataset” (30422 sequences). Donor-wise amino acid frequencies were computed for the sequences in the public clonotype the cAb dataset. The ratio of the per-residue amino acid frequencies was calculated with respect to the cAb dataset. The sequence logo plot to visualize residue-wise mutations and amino acid diversity was generated by using Weblogo (Crooks et al, 2004).

### Antigen expression and purification

BG505 gp140 SOSIP variants were expressed as recombinant soluble antigens. The single-chain variants (Georgiev et al., 2015) of BG505 and selected point mutants, each containing an Avi tag, were expressed in FreeStyle 293F mammalian cells (ThermoFisher) using polyethylenimine (PEI) transfection reagent and cultured for 5-7 days. FreeStyle 293F cells were maintained in FreeStyle 293F medium or FreeStyle F17 expression medium supplemented with 1% of 10% pluronic F-68 and 20% of 200 mM L-Glutamine. These cells were cultured at 37°C with 8% CO_2_ saturation and shaking. After transfection and 5-7 days of culture, cell cultures were centrifuged at 6000 rpm for 20 minutes. Supernatant was 0.45 µm filtered with PES membrane Nalgene Rapid Flow Disposable Filter Units and then run slowly over an affinity column of agarose bound Galanthus nivalis lectin (Vector Laboratories cat no. AL-1243-5) at 4°C. The column was washed with PBS, and proteins were eluted with 30 mL of 1 M methyl-α-D-mannopyranoside. The protein elution was buffer exchanged three times into PBS and concentrated using 30kDa Amicon Ultra centrifugal filter units. Concentrated protein was run on a Superose 6 Increase 10/300 GL or Superdex 200 Increase 10/300 GL sizing column on the AKTA FPLC system, and fractions were collected on an F9-R fraction collector. Fractions corresponding to correctly folded antigen were selected, and antigenicity by ELISA was characterized with known monoclonal antibodies specific for that antigen. Proteins were stored at –80°C until use.

For producing BG505 DS-SOSIP Env (Do Kwon et al., 2015) expressing high mannose glycans, we produced the Env in GnT1-cells. Env was affinity purified using a PGT145 IgG affinity column (De Taeye et al., 2015), followed by a Superdex 6 Increase 10/300 column.

### Antibody expression and purification

For each antibody, variable genes were inserted into plasmids encoding the constant region for the heavy chain (pFUSEss-CHIg-hG1, Invivogen) and light chain (pFUSE2ss-CLIg-hl2, Invivogen and pFUSE2ss-CLIg-hk, Invivogen) and synthesized from GenScript. mAbs were expressed in FreeStyle 293F or Expi293F mammalian cells (ThermoFisher) by co-transfecting heavy chain and light chain expressing plasmids using polyethylenimine (PEI) transfection reagent and cultured for 5-7 days. FreeStyle 293F (ThermoFisher) and Expi293F (ThermoFisher) cells were maintained in FreeStyle 293F medium or FreeStyle F17 expression medium supplemented with 1% of 10% pluronic F-68 and 20% of 200 mM L-Glutamine. These cells were cultured at 37°C with 8% CO_2_ saturation and shaking. After transfection and 5-7 days of culture, cell cultures were centrifuged at 6000 rpm for 20 minutes. Supernatant was 0.45 µm filtered with PES membrane Nalgene Rapid Flow Disposable Filter Units. Filtered supernatant was run over a column containing Protein A agarose resin that had been equilibrated with PBS. The column was washed with PBS, and then antibodies were eluted with 100 mM Glycine HCl at pH 2.7 directly into a 1:10 volume of 1 M Tris-HCl pH 8. Eluted antibodies were buffer exchanged into PBS three times using 10kDa or 30kDa Amicon Ultra centrifugal filter units.

### Enzyme linked immunosorbent assay (ELISA)

For gp120 ELISAs, soluble 93TH975 (Aids Reagent Program) protein was plated at 2 μg/ml overnight at 4°C. The next day, plates were washed three times with PBS supplemented with 0.05% Tween20 (PBS-T) and coated with 5% milk powder in PBS-T. Plates were incubated for one hour at room temperature and then washed three times with PBS-T. Primary antibodies were diluted in 1% milk in PBS-T, starting at 10 μg/ml with a serial 1:5 dilution and then added to the plate. The plates were incubated at room temperature for one hour and then washed three times in PBS-T. The secondary antibody, goat anti-human IgG conjugated to peroxidase, was added at 1:10,000 dilution in 1% milk in PBS-T to the plates, which were incubated for one hour at room temperature. Plates were washed three times with PBS-T and then developed by adding TMB substrate to each well. The plates were incubated at room temperature for 10 minutes, and then 1 N sulfuric acid was added to stop the reaction. Plates were read at 450 nm.

For recombinant single-chain SOSIP trimer ELISAs, 2 μg/ml of recombinant trimer proteins diluted in PBS were added to the plate and incubated overnight at 4°C. Primary and secondary antibodies, along with substrate and sulfuric acid, were added as described above.

Areas under the ELISA binding curves (AUC) were determined with GraphPad Prism 8.0.0.

### Surface Plasmon Resonance (SPR)

The binding of native CAP248_30 to different concentrations of BG505.DS.SOSIP/GnTI-was assessed by surface plasmon resonance on Biacore T-200 (Cytiva) at 25°C with HBS-EP+ (10 mM HEPES, pH 7.4, 150 mM NaCl, 3 mM EDTA, and 0.05% surfactant P-20) as the running buffer. 200nM of CAP248_30 IgG was captured on flow cell of immobilized human Anti-Fc chip (9,000 RU) and it was assayed by flowing over 50, 100, 200, 400nM of BG505.DS.SOSIP/GnTI-in running buffer. The surface was regenerated between injections by flowing over 3M MgCl2 solution for 10 s with flow rate of 100 µl/min. Blank sensorgrams were obtained by injection of the same volume of HBS-EP+ buffer in place of BG505.DS.SOSIP/GnTI-solutions. Sensorgrams of the concentration series were corrected with corresponding blank curves.

The binding of antibody CAP248_30_H_/CAP314_30_L_ to BG505 SOSIP/GnT1- and BG505.DS.SOSIP/GnT1-, and BG505.DS.SOSIP/293F (Do Kwon et al., 2015) was assessed by surface plasmon resonance on Biacore T-200 (GE-Healthcare) at 25°C with HBS-EP+ (10 mM HEPES, pH 7.4, 150 mM NaCl, 3 mM EDTA, and 0.05% surfactant P-20) as the running buffer. The antibody was captured on a CM5 chip by flowing 200 nM of the antibody over a flow cell immobilized with ∼9000 RU of anti-human Fc antibody. Binding was measured by flowing over 100, 200, 400 and 800 nM solution of BG505.DS.SOSIP/GnT1- and by flowing over 800 nM solution of BG505.SOSIP/GnT1- and BG505.DS SOSIP/293F in running buffer. The surface was regenerated between injections by flowing over 3M MgCl2 solution for 10 s with flow rate of 100 µl/min. Blank sensorgrams were obtained by injection of same volume of HBS-EP+ buffer in place of trimer solutions. Sensorgrams of the concentration series were corrected with corresponding blank curves.

### TZM-bl Neutralization Assays

Antibody neutralization was assessed using the TZM-bl assay as described (Montefiori, 2004; Sarzotti-Kelsoe et al., 2014). This standardized assay measures antibody-mediated inhibition of infection of JC53BL-13 cells (also known as TZM-bl cells) as a reduction in luciferase gene expression after a single round of infection by molecularly cloned Env-pseudoviruses. Murine leukemia virus (MLV) was included as an HIV-specificity control and VRC01 was used as a positive control. Titers (µg/ml) were calculated as the inhibitory concentration (IC_50_) causing 50% reduction of relative light units (RLU) with respect to the virus control wells (untreated virus). Reported titers represent an average of two or more repeat IC_50_ values with values <0.02 reported as 0.02 and values >50 reported as 50.

### Statistical Analysis

Spearman correlation tests were performed using cor.test function in R to obtain p and r values to determine the statistical significance of the differences. For the comparison of multiple independent tests, the p values were adjusted using the Benjamini-Hochberg method (Benjamini and Hochberg, 1995) using the p.adjust function in R. GraphPad Prism 8.0.0 was used to calculate 2 way ANOVAs as well as p-values adjusted for multiple comparisons via Tukey’s multiple comparison test.

