## Supplementary Figures for "Sequence and functional characterization of a public HIV-specific antibody clonotype"

**Supplementary Figure 1. Antibody Binding to (a) Monomer or (b) Trimer**

Antigen specificity of each antibody heavy-light chain pair was validated by ELISA against (a) 93TH975 gp120 monomer and (b) BG505.T332N.SOSIP.664 single-chain trimer. VRC01 or 3BNC117 were used as a positive control and are colored in black.

**Supplementary Figure 2**

**Public antibody binding to HIV-1 trimer by surface plasmon resonance.** Binding for (A) native CAP248_30 and (B) non-native CAP248_$30_H$/CAP314_$30_L$ antibodies was assessed against BG505.SOSIP.664 or a SOSIP variant with an additional disulfide stabilization (DS) (Do Kwon et al., 2015) expressed in 293F cells or GnT1- cells.

**Supplementary Figure 3. Antibodies from the Public Clonotype Map to the CD4 Binding Site of HIV-1 Env.**

AUC values from ELISA curves for a given antibody heavy-light chain pair against select epitope-specific point mutants of BG505.SOSIP.664.sc trimers as a percentage of binding to BG505.SOSIP.T332N.664.sc. Level of decrease of binding is shown as a heatmap. Antibodies with AUC values less than 0.5 are shown in grey or are absent. Known HIV-1 antibodies VRC01 and 10-1074 were used as controls. The public antibodies were susceptible to the double epitope knockouts D368R and D279K, suggesting targeting of the CD4 binding site.

**Supplementary Figure 4. Pseudovirus neutralization by public antibodies.** Neutralization $IC_{50}$ values for different antibody heavy-light chain pairs. $IC_{50}$ values are shown from high potency (< 0.1 µg/ml, dark red) to no neutralization (> 50 µg/ml, white).

**Supplementary Figure 5. Comparison of Residue-Specific (a) Somatic Hypermutation and (b) Entropy Between Antibodies from Different Donors in the Public Clonotype.** Each dot represents a single residue position in the sequence alignment for the public clonotype, showing the somatic hypermutation or entropy value for that residue position in each donor. Spearman correlations between all pairs of donors were computed and p-values corrected for multiple comparisons using the Benjamini-Hochberg method were reported. A total of 71 residue positions (from CDR1 to FR3) were plotted for the somatic hypermutation analysis and 85 residue positions (from CDR1 to CDR3) were plotted for the entropy analysis.

**Supplementary Figure 6. Donor-specific Amino Acid Frequency for Sequences in the Public Clonotype**

Heatmap representation of amino acid frequency for the sequences in the public clonotype (shown separately for each donor) and the cAb dataset. Residue positions from CDR1 to FR3 are shown in the heatmap. The frequency values ranged from 0 (white) to 1 (donor-based color).

### Supplementary Figure 1

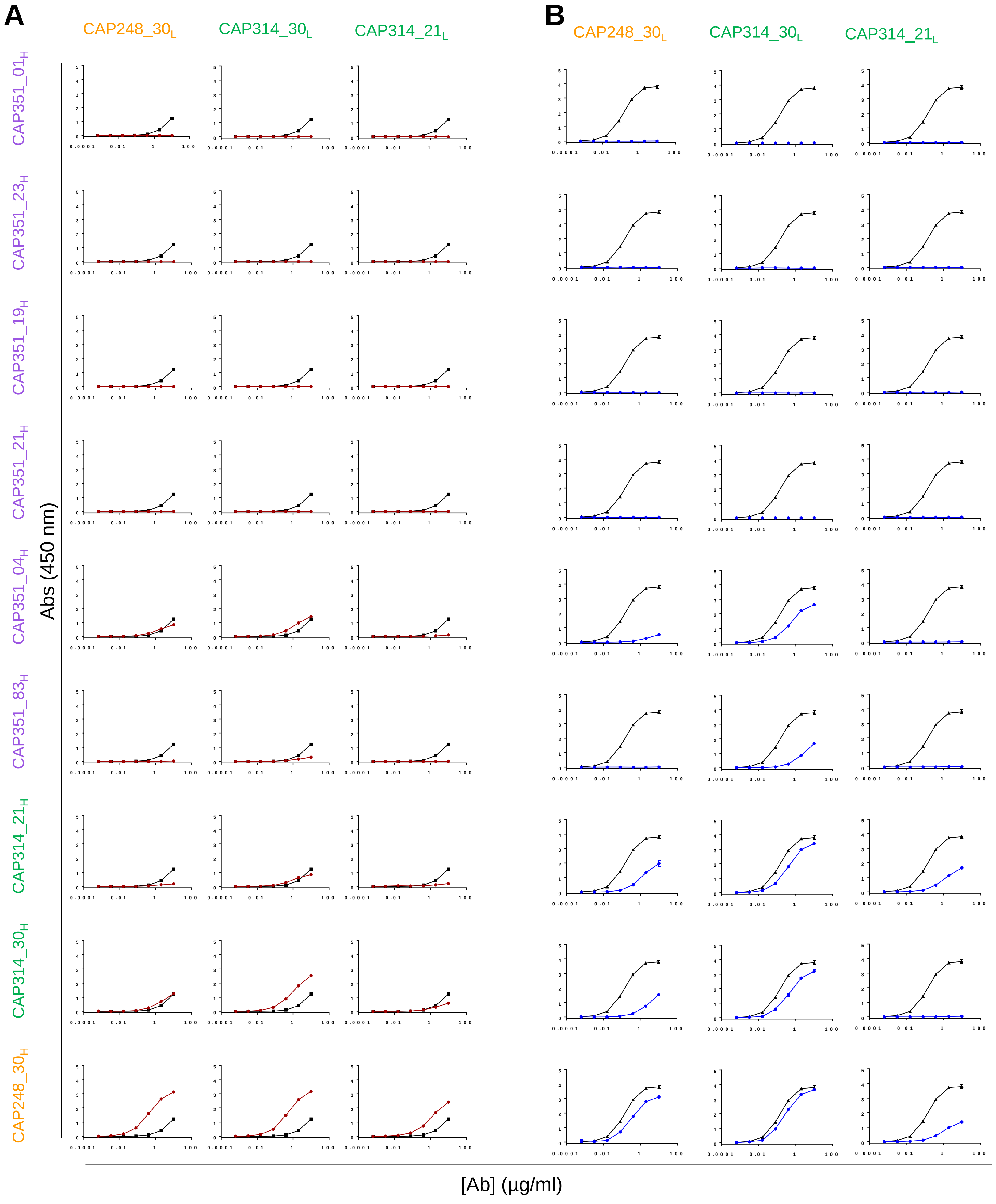

### Supplementary Figure 2

**A**

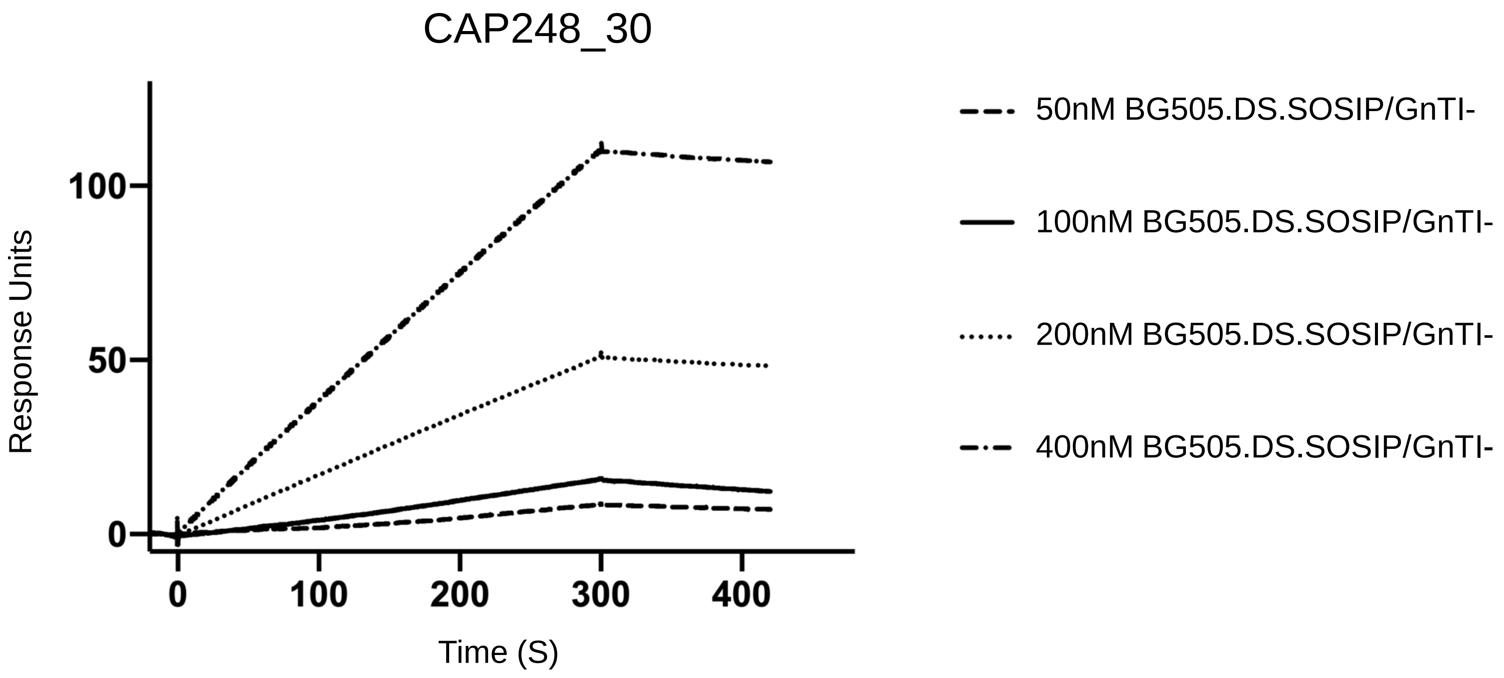


**B**

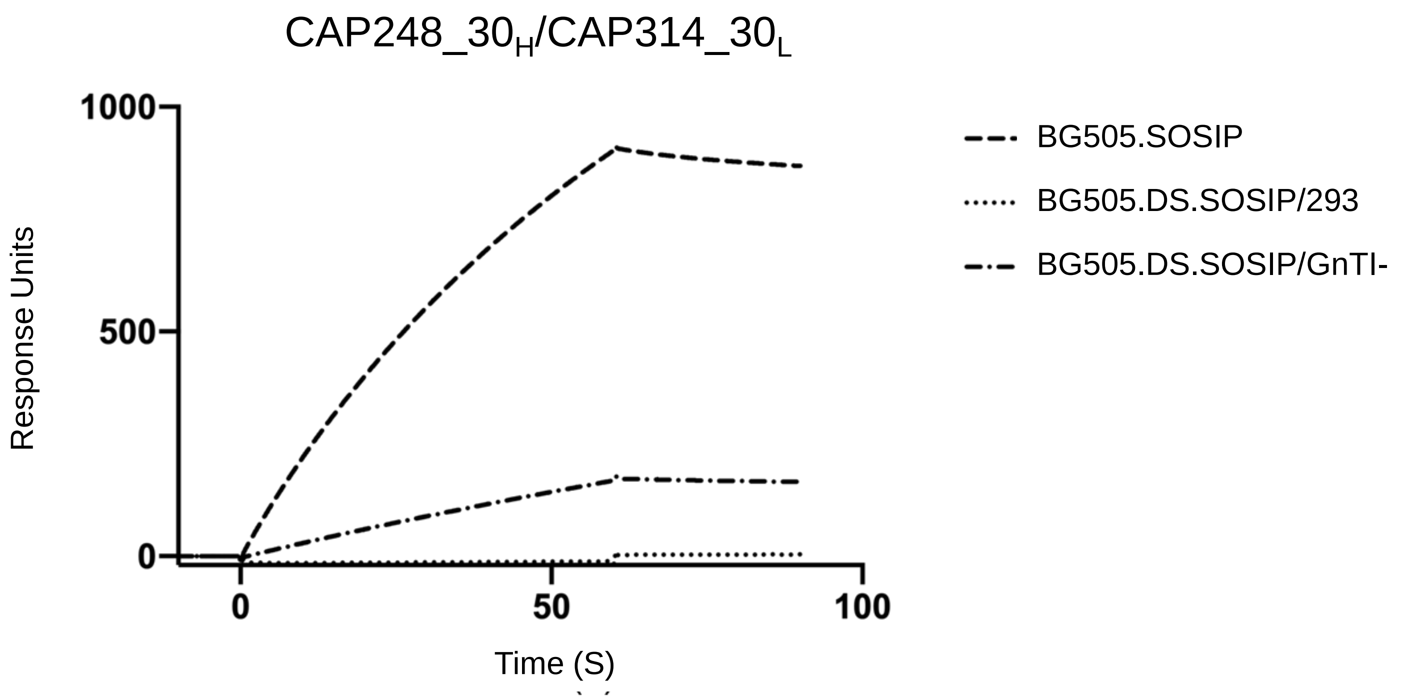

### Supplementary Figure 3

#### BG505.T332N.D279K.D368R.SOSIP.664.SC

| Heavy Chain \ Light Chain | CAP248_30 | CAP314_30 | CAP314_21 |
|---|---|---|---|
| CAP351_04 | | 0.00 | |
| CAP351_83 | | 0.03 | |
| CAP314_21 | 0.00 | 0.00 | 0.01 |
| CAP314_30 | 0.02 | 0.01 | |
| CAP248_30 | 0.00 | 0.00 | 0.00 |

| | |
|---|---|
| VRC01 | 0.00 |
| 10-1074 | 0.85 |

#### BG505.N332T.SOSIP.664.SC

| Heavy Chain \ Light Chain | CAP248_30 | CAP314_30 | CAP314_321 |
|---|---|---|---|
| CAP351_04 | | 1.06 | |
| CAP351_83 | | 1.08 | |
| CAP314_21 | 1.46 | 0.96 | 1.36 |
| CAP314_30 | 1.16 | 0.91 | |
| CAP248_30 | 0.89 | 0.87 | 1.58 |

| | |
|---|---|
| VRC01 | 1.01 |
| 10-1074 | 0.00 |

Biggest decrease in binding

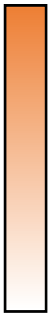

Smallest decrease in binding

### Supplementary Figure 4

MW965.26
Tier 1A, Clade C

| | CAP248_30LC | CAP314_30LC |
|---|---|---|
| CAP248_30HC | <0.02 | <0.02 |
| CAP314_30HC | 0.11 | 0.03 |

MN3
Tier 1A, Clade B

| | CAP248_30LC | CAP314_30LC |
|---|---|---|
| CAP248_30HC | 0.1 | 0.03 |
| CAP314_30HC | >50 | 0.2 |

6644
Tier 1B, Clade B

| | CAP248_30LC | CAP314_30LC |
|---|---|---|
| CAP248_30HC | 1.46 | 0.93 |
| CAP314_30HC | 4.34 | 3.12 |

| IC50 in µg/ml |
|---|
| <0.1 |
| 0.1-1.0 |
| 1.0-10.0 |
| 10.0-50.0 |
| >50 |

#### Supplementary Figure 5

**A**

Pairwise Comparisons of Somatic Hypermutation

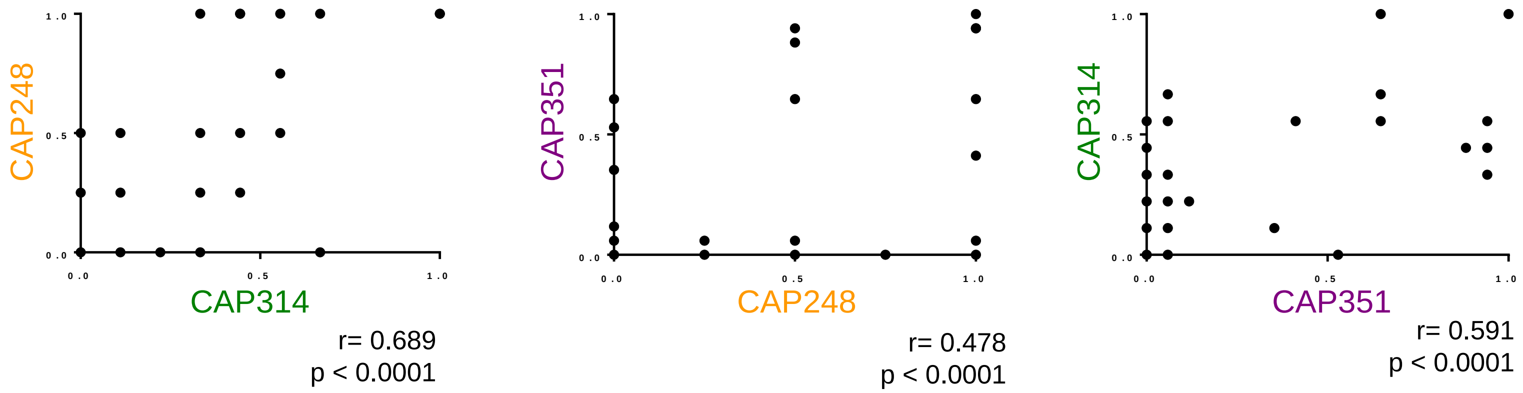


**B**

Pairwise Comparisons of Entropy Values

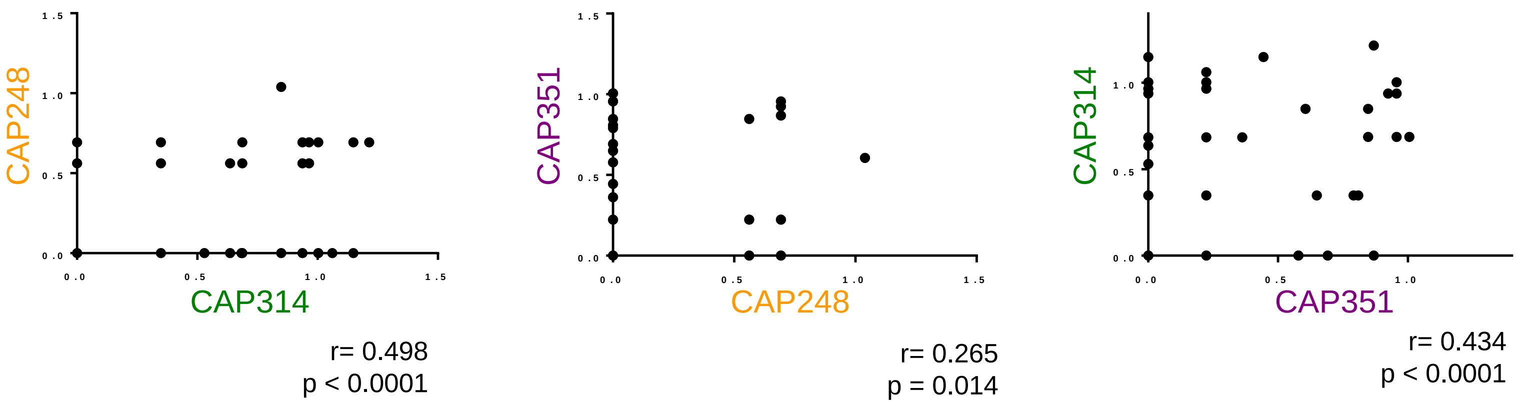

### Supplementary Figure 6

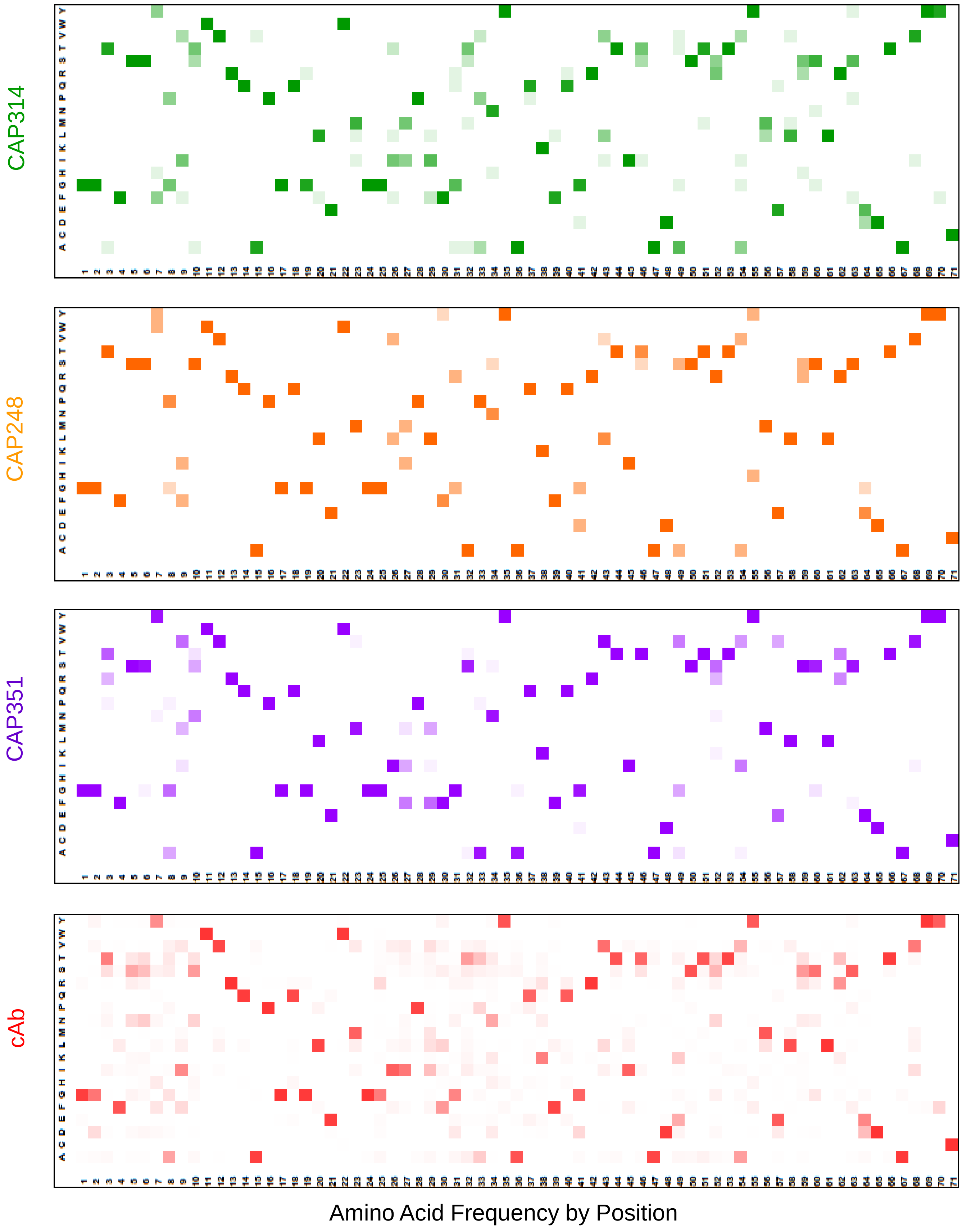
